# The role of task on the human brain’s responses to, and representation of, visual regularity defined by reflection and rotation

**DOI:** 10.1101/2023.10.27.564326

**Authors:** E Zamboni, A.D.J. Makin, M. Bertamini, A.B. Morland

## Abstract

Identifying and segmenting objects in an image is generally achieved effortlessly and is facilitated by the presence of symmetry: a principle of perceptual organisation used to interpret sensory inputs from the retina into meaningful representations. However, while imaging studies show evidence of symmetry selective responses across extrastriate visual areas in the human brain, whether symmetry is processed automatically is still under debate. We used functional Magnetic Resonance Imaging (fMRI) to study the response to and representation of two types of symmetry: reflection and rotation. Dot pattern stimuli were presented to 15 human participants (10 female) under stimulus-relevant (symmetry) and stimulus-irrelevant (luminance) task conditions. Our results show that symmetry-selective responses emerge from area V3 and extend throughout extrastriate visual areas. This response is largely maintained when participants engage in the stimulus irrelevant task, suggesting an automaticity to processing visual symmetry. Our multi-voxel pattern analysis (MVPA) results extend these findings by suggesting that not only spatial organisation of responses to symmetrical patterns can be distinguished from that of non-symmetrical (random) patterns, but also that representation of reflection and rotation symmetry can be differentiated in extrastriate and object-selective visual areas. Moreover, task demands did not affect the neural representation of the symmetry information. Intriguingly, our MVPA results show an interesting dissociation: representation of luminance (stimulus irrelevant feature) is maintained in visual cortex only when task relevant, while information of the spatial configuration of the stimuli is available across task conditions. This speaks in favour of the automaticity for processing perceptual organisation: extrastriate visual areas compute and represent global, spatial properties irrespective of the task at hand.

**Key Points:**

1. Symmetry selective responses observed in extrastriate visual cortex but not V1 whether stimulus spatial configuration is task relevant or irrelevant.
2. Representation for reflection and rotation differ in extrastriate and object-selective areas, during both stimulus relevant and irrelevant tasks.
3. Representation of luminance information is available only when task relevant.

## Introduction

Many animals, including humans, are sensitive to visual symmetry (Benard et al., 2006; Delius & Nowak, 1982; Treder, 2010; Wagemans, 1997). Symmetry can be found in both natural (e.g., faces and bodies, flowers, and crystals) and artificial (e.g., tools, buildings, and artworks) settings. An early observation about the role of symmetry in perceptual organisation comes from the Gestalt tradition. As one of the principles of grouping, symmetry helps in the process of segmenting and organising perceptual inputs into coherent and meaningful representations. Specifically, visual symmetry provides a strong cue for processes segregating a figure from its background (e.g., (Machilsen et al., 2009), making it an important component to shape representation (e.g., (Kovács & Julesz, 1993) and object recognition (Baylis & Driver, 1995; Bertamini et al., 1997).

Symmetry, or regularity more generally, has different forms and like other visual features (contrast; colour; (Anton-Erxleben & Carrasco, 2013; Di Russo et al., 2001; Morrone et al., 2004) research has investigated the extent to which those different forms of symmetry are processed in the context of attention. Reflectional symmetry has been suggested to be processed effortlessly and efficiently (Barlow & Reeves, 1979). However, the automaticity with which symmetry is processed and detected is still under debate: at one extreme this might occur pre-attentively (Wagemans, 1995), and at the other end of the spectrum this could be processed only when relevant to the task to be carried out (e.g., (Niimi et al., 2006; Wagemans, 1995). Recent behavioural work by (Kimchi et al., 2016), and (Devyatko & Kimchi, 2020) suggests that there are types of reflectional symmetry that are not processed pre-attentively. A perceptual difference between reflectional and rotational symmetry was already noted by Ernst Mach (1886). For humans, reflection is much more salient than rotation, even when these regularities are matched in terms of the number of isometric transformations (Royer, 1981; van der Helm & Leeuwenberg, 1996), and reflectional symmetry, not rotational symmetry, aids figure-ground segmentation (Machilsen et al., 2009) and is reinforced by closure (Bertamini, 2010). It might be that reflectional symmetry is special, and thus processed more readily than other types of regularity, such as rotational symmetry.

Neural signatures of symmetry processing in humans have been extensively studied with evoked potentials. The Sustained Posterior Negativity (SPN) is a regularity-driven, late component presenting a negative amplitude (Höfel & Jacobsen, 2007a; Makin et al., 2012, 2015, 2016; Norcia et al., 2002; Rampone et al., 2014), resulting from the difference between response to symmetrical and asymmetrical stimuli. Comparing neural responses to different types of symmetry (reflection, rotation, translation), results in a similar SPN response, which is therefore not unique to reflectional symmetry (Makin et al., 2013). This suggests an overlapping mechanism sensitive to different types of visual symmetry. However, the magnitude of the SPN is modulated by the type of symmetry (Makin et al., 2013), with the strongest response emerging for reflectional symmetry, followed by rotation and translation. The SPN has also been reported to be elicited automatically: it is present whether participants are attending to symmetry (i.e., this feature is task-relevant) or whether they are directing attention to other aspects of the visual stimuli (i.e., regularity is task-irrelevant), and even during passive viewing (i.e., no overt task required) (Bertamini & Makin, 2014; Höfel & Jacobsen, 2007a, 2007b; Makin, Rampone, Morris, et al., 2020) (Makin et al., 2022) reported analysis of all datasets from their complete Liverpool SPN catalogue (available on Open Science Framework, https://osf.io/2sncj/). They found that both stimulus and task (attention) manipulations influence SPN amplitude, with few tasks which completely abolish the SPN response to the most salient forms of regularity. While this field of research helps understanding the timing of visual symmetry processing, it lacks the spatial resolution to tease apart how this visual feature, together with its subgroups (e.g., reflection, rotation, translation) is specifically represented in the brain.

The localising of brain responses to and representation of symmetry has been investigated in animal models and humans. Regularity responses can be found in area V4 of anaesthetised monkeys (Audurier et al., 2022; Gallant et al., 1996), likely reflecting the important role for V4 in processing spatial information underpinning shape perception. However, regularity may be processed even earlier in the visual hierarchy as studies of the macaque visual system have shown regularity-specific activation in area V2 for visual stimuli capturing structural features of naturalistic textures ((Freeman et al., 2013). In human, primary visual cortex does not show a differential response when comparing activation to symmetrical versus random configurations (Chen et al., 2007; Keefe et al., 2018; Kohler et al., 2016; Sasaki et al., 2005; Tyler et al., 2005; Van Meel et al., 2019). (Sasaki et al., 2005) reports that symmetry related activation in visual areas V3A, V4, V7, and the Lateral Occipital Complex (LOC) parametrically changes with the signal-to-noise ratio present in the stimuli: the higher the percentage of symmetrical information available, the higher the measured activation in these extrastriate regions. More recently (Keefe et al., 2018), extended upon this idea by showing that the symmetry response in extrastriate areas is not only modulated by the amount of coherence in dot patterns, but also by the number of folds (axes of symmetry) present in the pattern. Importantly, whether observers were passively viewing the symmetrical dot patterns or engaging in a symmetry-detection task, resulted in robust symmetry-selective responses in these regions.

In this study we investigated response to and representations of two types of symmetry (reflection and rotation) under stimulus relevant and irrelevant task conditions. Specifically, we use two categories of visual symmetry: 4-fold reflections (or mirror symmetry), perceptually considered the most salient type of symmetry (Treder, 2010; Wagemans, 1997), and 45° rotational symmetry, together with random dot configurations. Attention was manipulated by requiring observers to perform a symmetry-relevant (i.e., press a key when two consecutive reflection/rotation/random stimuli appear) or symmetry-irrelevant (i.e., press a key when two consecutive patterns with the same luminance appear) task on the stimuli while patterns of activation from retinotopically identified visual areas were measured. We first undertook univariate analysis of the brain responses to compare with our work (Keefe et al., 2018) and that of others (Kohler et al., 2016). A recent study by (Van Meel et al., 2019) investigated dissimilarities in symmetrical and asymmetrical neural representations using classification analysis techniques. Here we took advantage of this technique to determine in which retinotopically defined regions, symmetrical patterns are encoded and whether these representations change under different attentional conditions. We hypothesise: (1) a difference in neuronal representations of symmetrical and random configurations will emerge in extrastriate visual areas; (2) representations of different symmetrical categories will be distinguishable in object-selective cortex; (3) different attentional conditions will modulate the neural representation of symmetrical patterns.

## Materials and Methods

### Participants

Fifteen participants were recruited to the study (mean age 27.8, SD 5; 10 female). Data from two participants were removed from the analysis as they did not complete all three sessions of the study, while two more datasets were removed due to at or below chance performance in the behavioural task. All participants had normal or corrected-to-normal visual acuity and no history of neurological impairments. All participants gave written informed consent prior to taking part in the procedure. The study was approved by the York Neuroimaging Centre (YNiC) Research Ethics Committee at the University of York (UK). Each participant underwent a minimum of two (3 hours in total) and maximum of three (4.5 hours in total) scanning sessions. The study comprised 60 hours of scanning in total.

### Stimuli

Stimuli, presented on a dark grey background, consisted of dot patterns generated using custom-based python scripts (available on Open Science Framework at https://osf.io/t7esz/?view_only=132bc7e6178741f4b2b10c045c355fcb). Specifically, each dot pattern consisted of 64 dots (0.016 degrees of visual angle in diameter), organised within a circular aperture spanning 4.18 dva. Example patterns are provided in Fig 1: each was generated by populating a quadrant of the pattern that was then reflected, resulting in a fourfold symmetry (reflection condition) or rotated (rotation condition). The random dot patterns were generated such that the number of elements into each quadrant was consistent with the regular patterns (reflection and rotation). The dots forming the patterns were also randomly assigned one of three luminance values such that a third of the trials had patterns with white dots, another third had patterns with light grey dots, and a last third had patterns with dark grey dots (Fig 1 and Fig2 B). This allowed us to have participants perform either a regularity task or a task judging the luminance of the dot patterns.

**Figure 1.**
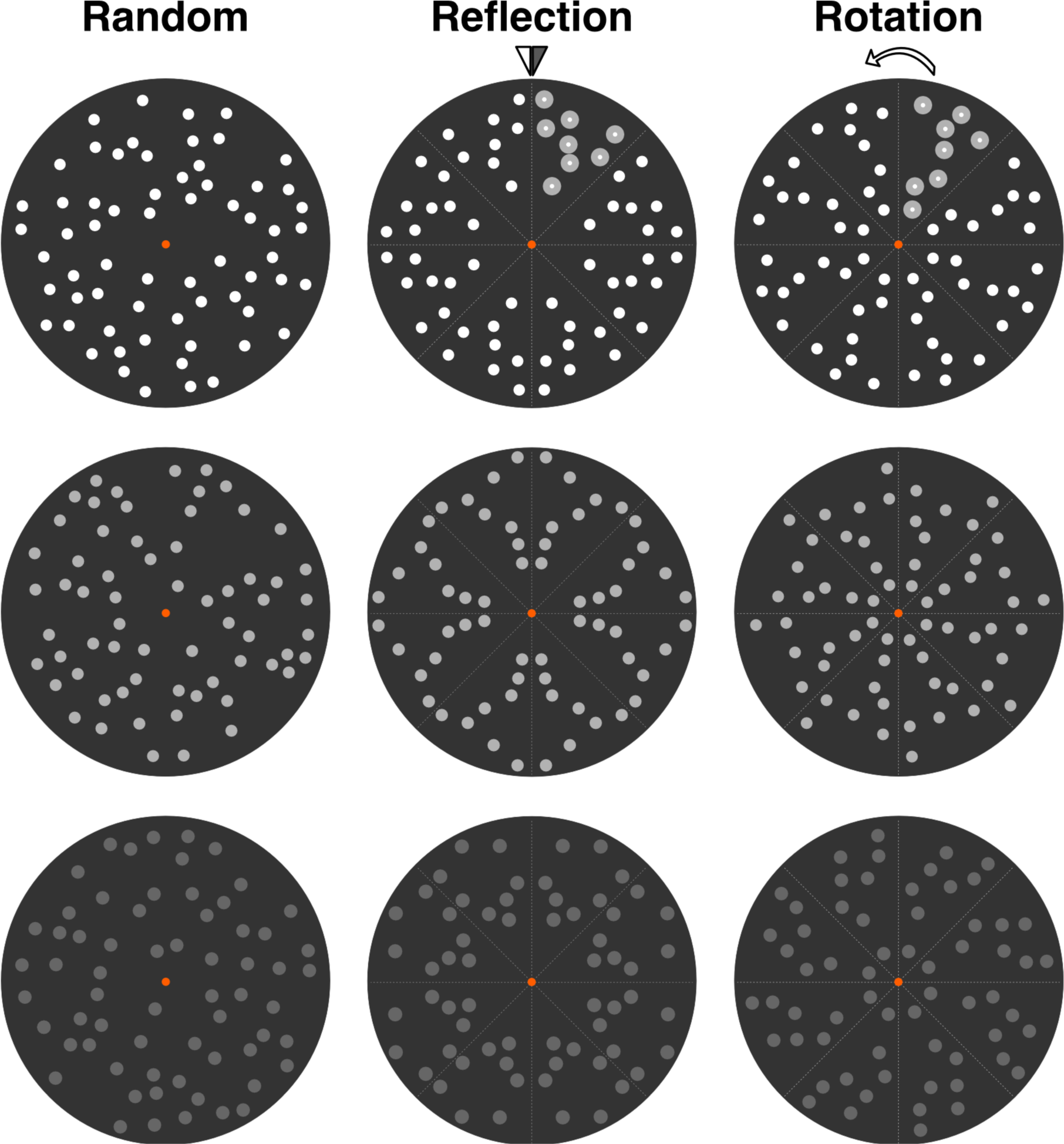
Example stimuli shown as circular apertures. Each column displays a different exemplar of a pattern type: random, reflection, rotation, respectively. Each row displays different levels of luminance used in the study. The segments with highlighted dots in the middle and right-most patterns (first row) indicate the kernels used to generate the rest of the patterns based on the geometrical transformation applied (i.e., 4-fold reflection or 45° rotation).

For each participant, seven unique patterns per condition (reflection, rotation, random) were generated during the first session and each pattern was presented four times throughout one run. Each individual participant was presented with the same patterns throughout the study, however patterns varied across participants, resulting in a large set of images onto which we base our Multi-Voxel Pattern Analysis (MVPA), largely reproducing the approach by (Van Meel et al., 2019).

The experiment was controlled using Python and open-source PsychoPy software (Peirce, 2007; Van Meel et al., 2019). Stimuli were presented using a projector and a mirror setup (1920 x 1080 pixels resolution, 120 Hz frame rate) at a viewing distance of 62 cm.

### Imaging Parameters

All imaging data were acquired on a Siemens MAGNETOM 3T Prisma scanner at the York Neuroimaging Centre (YNiC), University of York, using a 64-channel head coil for both functional and structural data. Each functional run consisted of T2*-weighted echoplanar images (EPIs) (52 slices, resolution 2.5mm isotropic, TR = 1000ms, TE = 30ms, 80x80 acquisition matrix, Multi-Band factor = 4, flip angle = 75 deg).

A T1-weighted high resolution anatomical scan was collected for each participant during the first session (320 slices, resolution 0.8mm isotropic, TR = 2400ms, TE = 2.28ms, FOV = 256x256, flip angle = 8 deg). To aid alignment of functional data to the T1-weighted, high resolution anatomical image, a Turbo Spin-Echo image, with the same prescription as the functional data was acquired (52 slices, resolution 1x1x2.5mm, TR = 7270ms, TE = 9.2ms, FOV = 200x200, flip angle = 160 deg).

### Experimental Procedure

The study consisted of two, 1.5-hour sessions, plus a third, 1-hour session for retinotopic mapping and LOC functional localiser. Two participants only completed one experimental session and were therefore removed from the analysis. For five participants, retinotopic mapping and functional localiser were already available from previous studies, the remaining eight participants completed all three sessions. Over the course of the two experimental sessions, participants performed a total of 10 runs (5 per task, in counterbalanced order, each lasting 7 min 15 s); during the third session, participants performed between 1 and 3 LOC functional localisers (each lasting 6 min 20 s; see (Vernon et al., 2016)), and between 4 and 8 retinotopic mapping scans (each lasting 2 min 8 s).

### Dot array experiments

For the experimental sessions, we used a rapid-event related design, in which the order of stimulus presentation was counterbalanced and optimised using Optseq2 (https://surfer.nmr.mgh.harvard.edu/optseq). Five stimulus presentation sequences were generated per participant, per session, such that a total of 84 events were presented per run (21 unique stimuli, each repeated four times). Stimuli were presented centrally for 1s with a jittered inter-stimulus interval (ISI) of between 3 and 11 s. Each run started with 10s and ended with 25s of fixation to ensure we captured the complete haemodynamic response for the last stimulus (see (Vernon et al., 2016). Participants maintained fixation on an orange central dot throughout the run and performed a one-back task on (1) the regularity of the patterns (press a button whenever two consecutive reflection / rotation / random patterns are presented) and (2) the luminance of the patterns (press a button whenever two consecutive white / light grey / dark grey patterns are presented). Participants were made aware of the task to be performed prior to entering the scanner and instructions were displayed on screen before the start of each run. During each scan, we recorded video of the participant’s left eye and later used these recordings to extract eyeblinks using custom-written software (see (Vernon et al., 2016).

### Retinotopic Mapping

For the retinotopic mapping session, we followed procedures previously described (see (Keefe et al., 2018; Welbourne et al., 2018). Briefly, a bar stimulus (width 0.5 deg) moved in one of eight possible directions within a 10 deg radius circular aperture. A movement across the full field lasted 16s, followed by a movement across half the direction for 8s, and interleaved with a mean luminance blank period for 8s before starting the cycle again. Participants performed a minimum of 4 and maximum of 8 scans.

### Functional Localiser

For the LOC functional localiser scan, we followed procedures previously described (see (Vernon et al., 2016). Briefly, an ABAB block design was used to present images of objects and scrambled objects, for a total of 16 blocks per condition. Each image was on-screen for 0.8s followed by 0.2s inter-stimulus interval; participants were asked to fixate a central red cross and to perform a one-back task on the images presented.

## Data Analyses

### Preprocessing and General Linear Model

T1-weighted structural data was used for coregistration and 3D cortex reconstruction. Grey and white matter segmentation for each hemisphere was obtained using Freesurfer (https://surfer.nmr.mgh.harvard.edu/) and further manually edited when necessary using ITKSnap (www.itksnap.org).

All functional data from the main experiment were analysed using FSL FEAT (fMRI Expert Analysis Tool; (Jezzard et al., 2001). Preprocessing steps consisted of distortion correction due to non-zero off-resonance field: at the beginning of each functional run, one volume with inverted phase encoding direction was acquired and used to estimate a voxel displacement map, subsequently applied to the functional volumes using FSL topup (Andersson et al., 2003; Smith et al., 2004). Once undistorted, the functional images underwent slice timing correction and high-pass filtering (90s cut-off point) to remove low-frequency drift. Functional volumes were motion corrected using MCFLIRT and co-registered to each individual’s structural image using boundary-based registration tools.

Following preprocessing, a general linear model was applied to the functional images at an individual (first) level, for each task dataset (Luminance, Regularity) separately:

#### (1)#Univariate analysis

The model consisted of 3 regressors of interest (i.e., one per stimulus category: reflection, rotation, random), and six regressors for the motion correction parameters as covariates of no experimental interest. Moreover, a fourth regressor was added to the model accounting for blinks during stimulus presentation, as they have been reported to be potential sources of noise (Gouws et al., 2014; Hupé et al., 2012). Contrasts were set up to compare each stimulus category against baseline. To combine data within a participant, we ran fixed effects analysis with cluster correction (Z>2.3, *p*<0.05). Mean percent signal change was then computed by visual area (see Regions of Interest and Functional Localiser section) using FeatQuery.

#### (2)#MVPA analysis

To extract information regarding the *regularity* of the stimuli, the model included 21 regressors of interest (i.e., one per stimulus type: 7 reflection, 7 rotation, 7 random), plus one modelling eye blinks, six regressors for the motion correction parameters, and one extra covariate modelling change in dot luminance.

To extract information regarding the *luminance* of the stimuli, the model included 3 regressors of interest (i.e., one per luminance level: white, light grey, dark grey), plus one modelling eye blinks and six regressors for the motion correction parameters.

For both analyses, contrasts were set up to compare each stimulus category to baseline. Data were not combined within participants and resulting contrasts (z-statistics) from each run were used in the training and testing routine of the MVPA analysis.

### Regions of Interest and Functional Localiser

Retinotopy data were processed using mrVista analysis software (https://web.stanford.edu/group/vista/cgi-bin/wiki/index.php/Software) (Vista Lab, Stanford University): within run head motion was corrected and functional runs were aligned to the high-resolution structural image using, as intermediate step, a Turbo Spin-Echo image acquired with the same prescription as the functional data. Aligned functional data were then averaged and correlational analysis followed standard procedures (e.g., (Keefe et al., 2018; Welbourne et al., 2018). The resulting phase maps were visualised onto flat patches centred around the occipital pole and used to identify the following regions of interest (ROIs) in both hemispheres: V1, V2, V3, V4, VO1, VO2, LO1, and LO2.

Data from the LOC functional localiser runs were processed and analysed following procedures described in (Vernon et al., 2016). Briefly, data were slice-time corrected, high pass filtered (90s cut-off), and motion corrected using MCFLIRT. No spatial smoothing was applied, while FILM prewhitening was used. Data was combined, where possible, within participants using fixed effects analysis with cluster correction (Z>2.3, *p*<0.05). The resulting maps were then used, at the individual level, to guide identification of LO1 and LO2 boundaries whenever data from the retinotopic maps alone were not sufficient. ROIs were further restricted by eccentricity (stimulus size) based on a stimulus vs fixation contrast obtained from the main experiment functional data (across sessions).

### Multi Voxel Pattern Analysis

We used multi voxel pattern analysis (MVPA) to investigate the representations of different regular patterns (i.e., reflection and rotation) across the visual cortex. We trained a linear Support Vector Machines (SVM, (Schölkopf et al., 1999) classifier using LIBSVM (https://www.csie.ntu.edu/~cjlin/libsvm/; (Chang & Lin, 2011; Vernon et al., 2016) implemented in MATLAB to discriminate: (a) reflection from random patterns; (b) rotation from random patterns; and (c) reflection from rotation patterns, when participants were attending to the *regularity* of the patterns vs when they were attending to the *luminance* of the patterns. Classification accuracy for each condition was computed using a leave-one-run-out cross-validation (Kriegeskorte et al., 2009): data for each participant, for each task, and each ROI were divided into training and test datasets, with a minimum of 28 training patterns (n=1 participant with 3 runs per task) and a maximum of 70 training patterns (for n=1 participant with 6 runs per task; the remaining participants had 5 runs per task, hence 56 training patterns) and 14 patterns for the test set. This method of cross-validation is important to limit overfitting the classifier (Valente et al., 2021): by iteratively generating a model on different training-testing sets, the potential of an extremely good/bad fit driving the overall performance is attenuated (Tong & Pratte, 2012); Cawley & Talbot, 2010). Classification accuracy was then averaged across validation sets, for each condition, and across participants, for each ROI and task, separately. The resulting accuracies were then tested against chance performance using one-sample tests (Kriegeskorte et al., 2007; Wailes-Newson et al., 2019) for each ROI separately, with ɑ-level Bonferroni corrected for effects of stimulus category (the higher the classification accuracy, the more distinct the representations of different regular patterns are for a specific area in the visual cortex). Performance of each independent ROI classifier is measured by percent correct classification (accuracy), with the null hypothesis of the classifier to perform at chance level (e.g., 50% for a two-way classification routine). Univariate t-test therefore allows to test whether classification accuracy is significantly better than chance, and a significant t-value indicates that the region’s response contains information about the experimental condition (Kriegeskorte et al., 2006).

## Results

### Univariate Responses

We identified visual areas using standard retinotopic mapping procedures as shown in Figure 2A for one representative participant and measured the BOLD response to dot patterns that were either random, symmetrical (4-fold reflection), or rotational (exemplars depicted in Figure 2B, from left to right, respectively). Statistical maps in response to each individual stimulus category for a representative participant are provided in Figure 2C (random>baseline, reflection>random, rotation>random, respectively). This standard contrast approach appears to show more widespread responses to reflection vs random compared to rotation vs random. However, the ROI analysis we employ offers a more sensitive way of assessing stimulus selectivity as presented below.

**Figure 2.**
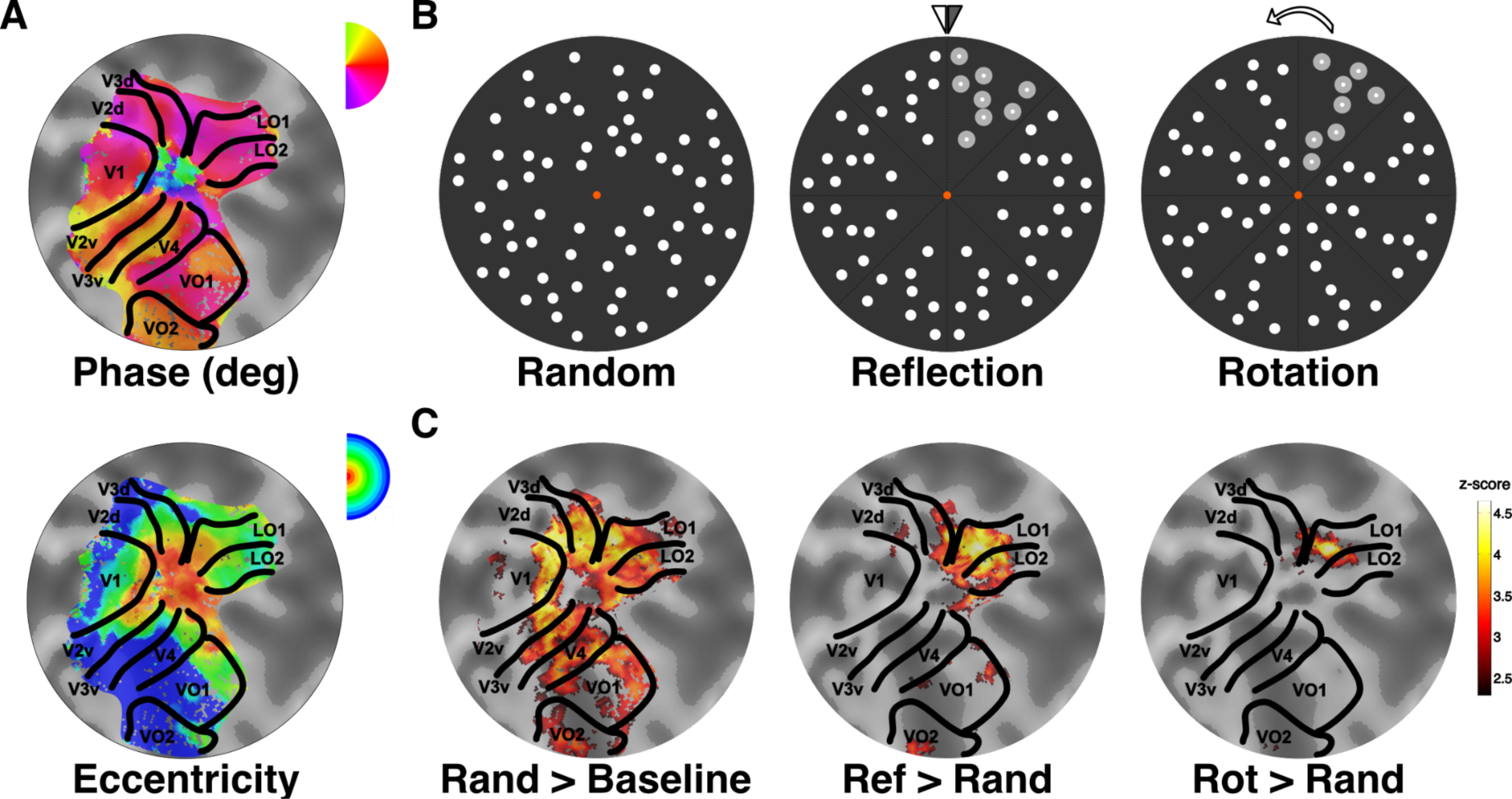
(A) Retinotopically defined ROIs: flat patch obtained from the occipital pole overlaid with a colour map detailing polar angle data (top) and eccentricity data (bottom). Visual areas are then drawn and labelled according to the reversals in phase (top) that demarcate them. (B) Example stimuli shown as circular apertures for random, reflection, and rotation, respectively. The segments with darker dots in the middle and right-most patterns indicate the kernels used to generate the rest of the patterns based on the geometrical transformation applied (i.e., 4-fold reflection or 45° rotation). (C) Heat maps indicating response to each stimulus category: random, reflection, and rotation. Note the symmetry-selective responses in extrastriate areas when contrasting response to reflection (middle) to random (left) and rotation (right) to random (left), while early visual areas V1 and V2 show no changes across categories (no symmetry-selective response observed in the middle and right maps). Data shown here are for a single participant, z-score thresholding as indicated by the colour bar insert.

The responses to the reflection, rotation and random dot stimuli are presented in Figure 3. Data are separately plotted for the two task conditions of luminance and regularity in panels A and B, respectively, and then across both tasks in panel C. Data are presented this way to allow visualisation of the effects that emerged from the statistical analysis presented below. It appears that there is little or no preferential response to symmetry in primary visual cortex and symmetry selectivity emerges more clearly up the visual hierarchy.

**Figure 3.**
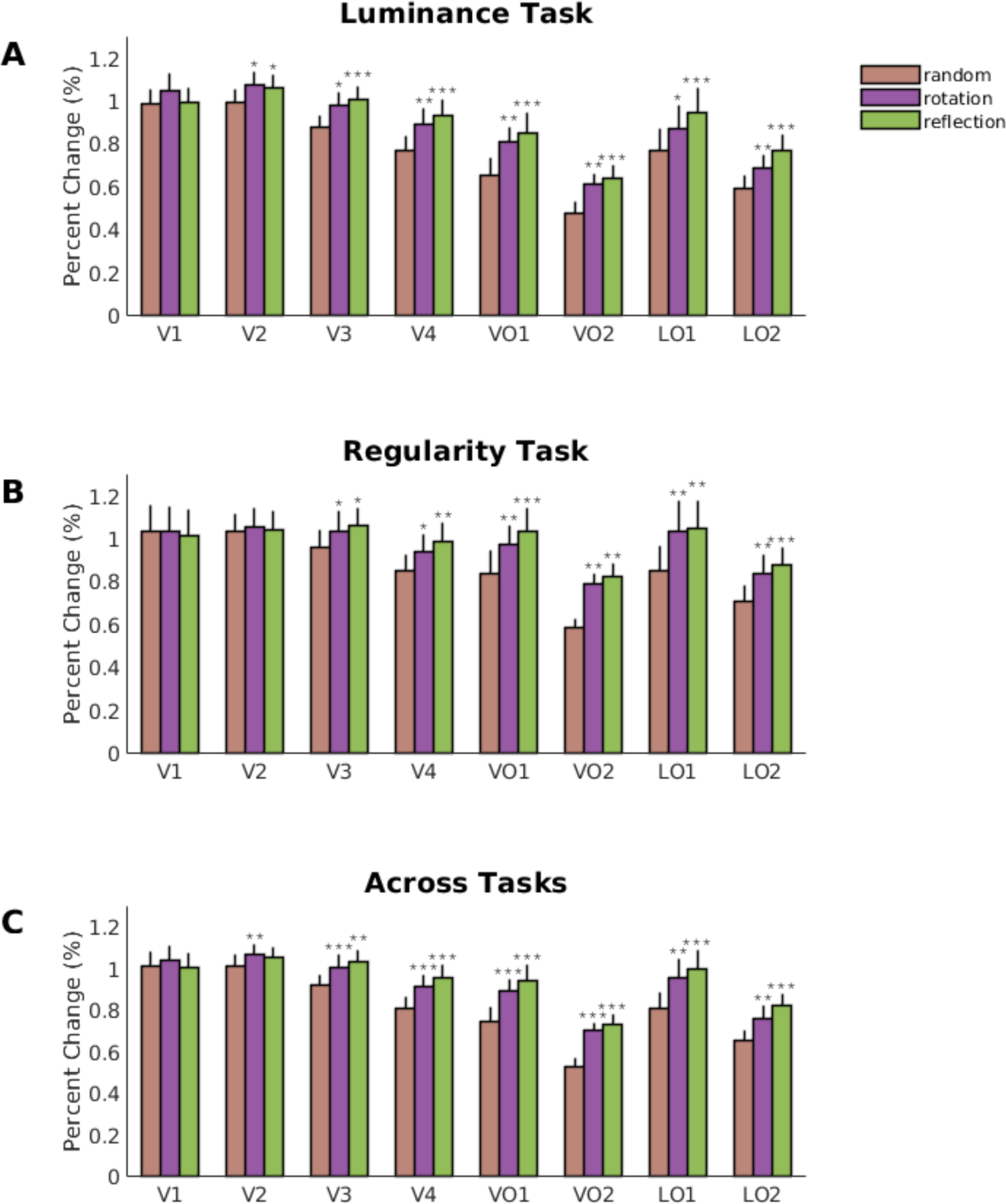
Univariate results for (A) luminance, (B) regularity, and (C) across tasks. Visual areas are represented along the x-axis, percent signal change is shown on the y-axis. Responses to random patterns are in light brown, responses to rotational patterns in purple, responses to reflection in green. Error bars indicate +SEM. Asterisks indicate areas where both responses to rotation and reflection were significantly higher than responses to random patterns: *<0.05, **<0.01,***<0.001 consistent with statistics reported in Table 1.

We investigated whether attending to regularity or luminance modulated the BOLD response across the visual cortex using a three-way, repeated measures ANOVA with factors of Task (attending to luminance, regularity), ROIs (V1, V2, V3, V4, VO1, VO2, LO1, LO2) and Stimulus (random, reflection, rotation). The degrees of freedom for the factor ROI were corrected using Huynh-Feldt (estimated epsilon larger than .5) following a departure from sphericity. There was a significant three-way interaction (F(6.546,64.560)=2.507, *p*=0.027, η^2^=0.2) showing that the way in which responses to the stimuli vary by ROI is dependent on the task. An indication of this is evident from comparing Figs 3A and B, where the regularity task appears to boost differences in responses to stimuli in extrastriate regions.

The application of the 3-way ANOVA also highlighted one significant two-way interaction; ROI by Stimulus (F(8,856,88.558)=10.886, *p*<0.0001, η^2^=0.521) and is illustrated in Fig 3C, where it is clear that symmetry selective responses emerge in extrastriate cortex. The other two-way interactions that depended on task did not reach significance; Task by ROIs (*p*=0.066) and Task by Stimulus (*p*=0.988). Main effects of ROI and Stimulus, but not Task were also significant; (ROI: F(3.681,36.830)=5.386, *p*=0.002, η^2^=0.35; Stimulus: F(2,20)=12.566, *p*=0.001, η^2^=0.557, respectively), Task: F(1,10)=3.484, *p*=0.092, η^2^=0.258)

To follow up on the three-way interactions we employed two-way ANOVAs with Stimulus and Task as factors for each ROI (each test and its post host, pairwise t-tests are given in Table 1). The analyses revealed a main effect of Stimulus across regions beyond V1 and V2, with areas V3, V4, VO1, VO2, LO1, and LO2 all showing an enhanced response to regular patterns (reflection and rotation) compared to random configurations, irrespective of task (all *p*<0.001). Interestingly, we also observed an effect of Task in ventral visual areas VO1 and VO2, where attending to the regularity of the patterns resulted in a larger BOLD response compared that when attending to the luminance of the stimuli (F(1,10)=11.87, *p*=0.006, η^2^=0.534; F(1,10)=5.915, *p*=0.035, η^2^=0.372 for VO1 and VO2, respectively). It appears therefore that the three-way interaction is driven by Task significantly boosting responses in VO1 and VO2, but not in other extrastriate areas, which do exhibit stimulus, but not task related responses.

**Table 1.** Results of two-way ANOVA performed on each region of interest. ROIs are shown in each row, while columns show data for the interaction (Task x Stimulus), main effects of Task (luminance, regularity), and Stimulus (random, reflection, rotation). Post-hoc t-tests (Task, Reflection vs Random, Rotation vs Random, Reflection vs Rotation) assess differences in task and stimulus types.

| ROI | Task x Stim<br>(F, p, $\eta^2$) | Task<br>(F, p, $\eta^2$) | Stim<br>(F, p, $\eta^2$) | Task<br>(t, p) | Reflection<br>vs Random<br>(t, p) | Rotation<br>vs Random<br>(t, p) | Reflection<br>vs Rotation<br>(t, p) |
| --- | --- | --- | --- | --- | --- | --- | --- |
| V1 | .444, .609,<br>.043 | .060, .811,<br>.006 | .891, .396,<br>.082 | .24,<br>.811 | -.237, 1.000 | 1.937, .239 | -1.111, .877 |
| V2 | .958, .396,<br>.087 | .000, .993,<br>.000 | 2.242,<br>.154, .183 | .015,<br>.993 | 1.500, .481 | **3.571, .012** | -.343, 1.000 |
| V3 | .217, .807,<br>.021 | 1.598,<br>.235, .138 | **11.074,**<br>**.001, .525** | 1.271,<br>.235 | **4.259, .005** | **4.684, .003** | .896, 1.000 |
| V4 | .209, .813,<br>.020 | 1.401,<br>.264, .123 | **14.958,**<br>**<.001, .599** | 1.196,<br>.264 | **5.000, .001** | **5.047, .001** | 1.333, .618 |
| VO1 | .038, .963,<br>.004 | **11.870,**<br>**.006, .543** | **17.934,**<br>**<.001, .642** | **3.471,**<br>**.006** | **6.758, <.0001** | **4.677, .002** | 1.243, .730 |
| VO2 | .881, .430,<br>.081 | **5.915,**<br>**.035, .372** | **16.138,**<br>**<.001, .617** | **2.406,**<br>**.035** | **5.075, .001** | **4.857, .002** | .825, 1.000 |
| LO1 | 1.507, .246,<br>.131 | 3.719,<br>.083, .271 | **13.730,**<br>**<.001, .579** | 1.931,<br>.083 | **5.108, .001** | **4.000, .007** | 1.125, .861 |
| LO2 | .474, .630,<br>.045 | 4.674,<br>.056, .319 | **13.179,**<br>**<.001, .569** | 2.155,<br>.056 | **5.581, .001** | **3.700, .013** | 1.550, .463 |

In summary, our univariate analysis showed: (1) an extrastriate network of visual areas starting beyond V2 that process regularity; this is in line with our hypothesis and previous literature, supporting the reliability of this result; (2) regularity is processed automatically, whether we are attending to the configuration (i.e., regularity-relevant), or to local features (i.e., regularity-irrelevant, here luminance) of the patterns, with the exception of more ventral areas that show also a larger response when attending to regularity-relevant features; (3) the extrastriate network exhibits indistinguishable univariate response to the two types of regularity; reflection and rotation.

### Multivariate Responses

The advantage of our design is that it allows us to exploit not only the average response of an ROI to a set of stimuli, but also look at the spatial organisation of the responses and attempt to discriminate between the response patterns relating to one stimulus category compared to another. To this end, we used multivoxel-pattern analysis techniques and trained an SVM classifier to distinguish between: (a) reflection and random patterns; (b) rotation and random patterns. We then run a one-sample t-test for each task condition (attending to luminance, attending to regularity) decoding accuracy against chance level (i.e., 50% accuracy of distinguishing between representation of the regular vs random configurations) separately for each ROI. The univariate analysis reported above already showed a significant difference between the mean BOLD response for these conditions. We would therefore expect our MVPA results to largely reflect the univariate findings.

Decoding accuracy for reflection vs random classification (Figure 4A) showed significantly above chance results for both attentional manipulations (luminance and regularity tasks), for all ROIs except V1 (V1: t(10)=1.994, *p*=0.074 and t(10)=2.068, *p*=0.065 for regularity and luminance tasks, respectively; all other ROIs show *p*<0.039 for both regularity and luminance tasks). Except for V2, the results are in line with the univariate analysis reported in the previous section.

**Figure 4.**
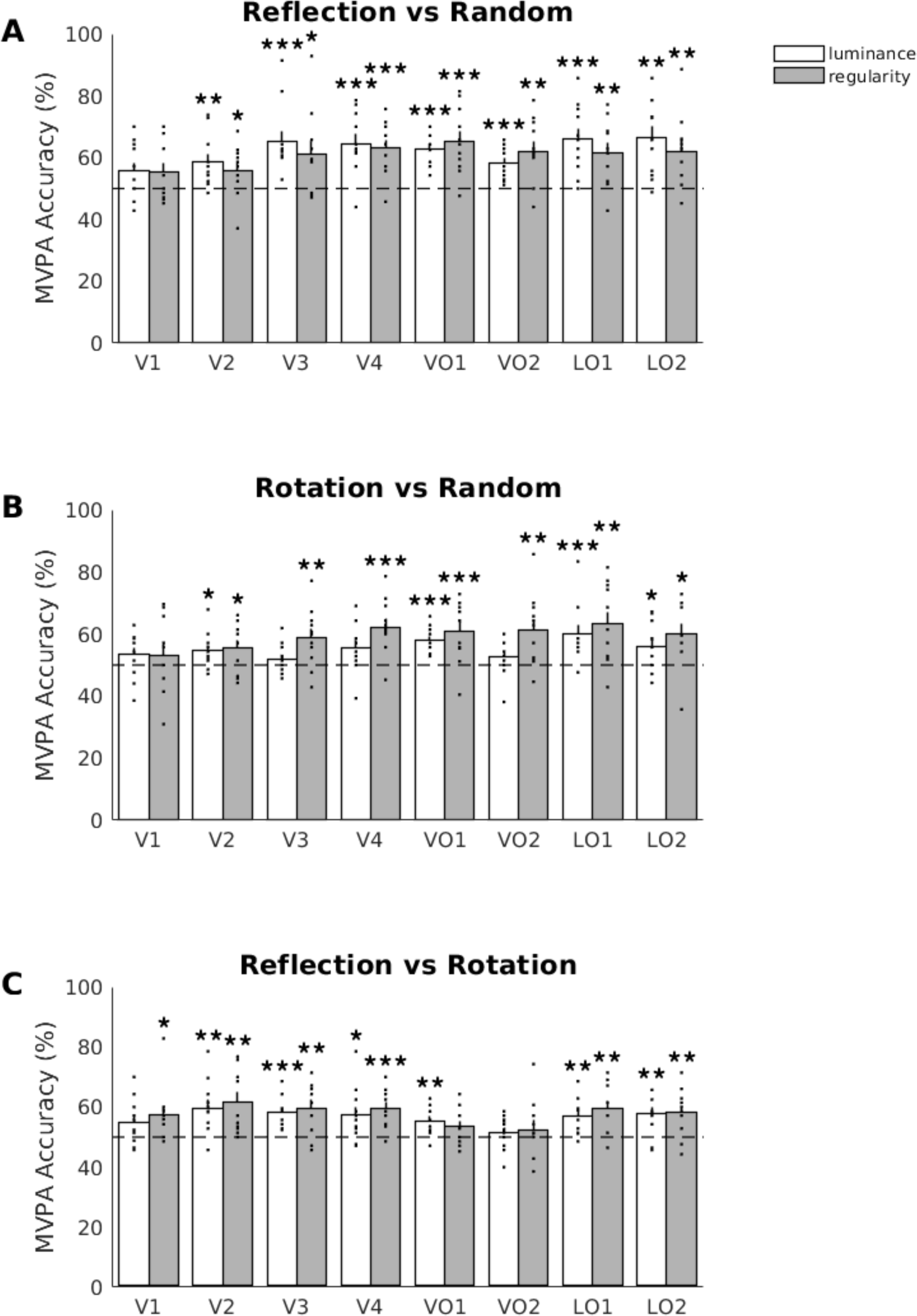
MVPA results for different stimulus conditions across visual ROIs. (A) Reflection vs Random: extrastriate areas show above chance classification accuracy during both the luminance (white) and the regularity (grey) tasks. (B) Rotation vs Random: extrastriate areas show above chance classification accuracy during regularity task, while attending to local features (i.e., luminance of dot patterns) results in above chance classification in ventral areas higher up the hierarchy (VO2, LO1, and LO2). (C) Reflection vs Rotation: extrastriate areas show above chance classification during both luminance and regularity tasks, with the exception of ventral areas VO1 and VO2. Individual dot markers indicate single subject data. Dashed lines indicate MVPA accuracy at 50% chance. Error bars indicate +SEM. Asterisks indicate significance as follows: *<0.05, **<0.01, ***<0.001 consistent with statistics reported in Table 2.

One sample t-test on decoding accuracy for rotation vs random condition (Figure 4B; results for each t-test performed, including estimate of effect size based on Hedge’s g computation to account for small sample size reported in Table 2) showed above chance classification accuracy when attending to regularity for all extrastriate ROIs, including V2 (*p*<0.0353). When attending to luminance, extrastriate areas VO1, LO1, and LO2 (p<0.031) still show above chance classification accuracy, as well as area V2 (*p*>0.052 for the other ROIs). These results suggest that rotation can be decoded at rates significantly above chance early in the visual hierarchy when attending to regularity.

**Table 2.** Results of one-sample t-tests assessing classification accuracy against 50% chance level for each stimulus pairing and task, across regions of interest. ROIs are given in each row and columns show statistical t, p-value, and Hedge’s g as a measure of effect size accounting for small sample size, for classification test (reflection vs random, rotation vs random, reflection vs rotation) and each task (luminance, regularity).

|  | Reflection vs Random<br>(t, p, Hedge's g) |  | Rotation vs Random<br>(t, p, Hedge's g) |  | Reflection vs Rotation<br>(t, p, Hedge's g) |  |
| --- | --- | --- | --- | --- | --- | --- |
| ROI | *luminance* | *regularity* | *luminance* | *regularity* | *luminance* | *regularity* |
| V1 | 2.068, .065, .575 | 1.994, .074, .555 | 1.499, .165, .417 | .887, .396, .247 | 2.127, .059, .592 | **2.679, .023, .745** |
| V2 | **3.476, .006, .967** | **2.367, .040, .658** | **2.654, .024, .738** | **2.433, .035, .677** | **3.617, .005, 1.006** | **3.821, .003, 1.063** |
| V3 | **4.568, .001, 1.271** | **2.823, .018, .786** | 1.105, .295, .307 | **3.127, .011, .870** | **4.901, .001, 1.364** | **3.769, .004, 1.049** |
| V4 | 4.720, .001,<br>1.313 | 4.967, .001,<br>1.382 | 2.208, .052,<br>.614 | 4.596, .001,<br>1.279 | 2.692, .023,<br>.749 | 5.075, <.001,<br>1.412 |
| VO1 | 8.145, <.001,<br>2.266 | 4.787, .001,<br>1.332 | 6.337, <.001,<br>1.763 | 3.701, .004,<br>1.030 | 3.326, .008,<br>.925 | 2.006, .073,<br>.558 |
| VO2 | 5.180, <.001,<br>1.441 | 3.682, .004,<br>1.024 | 1.373, .200,<br>.382 | 3.241, .009,<br>.902 | .961, .359,<br>.267 | .760, .465,<br>.212 |
| LO1 | 4.820, .001,<br>1.341 | 3.785, .004,<br>1.053 | 3.405, .007,<br>.947 | 3.425, .006,<br>.953 | 3.666, .004,<br>.1020 | 3.934, .003,<br>1.095 |
| LO2 | 4.376, .001,<br>1.217 | 3.426, .006,<br>.953 | 2.497, .032,<br>.695 | 3.148, .010,<br>.876 | 3.672, .004,<br>.1021 | 3.455, .006,<br>.961 |

While the previous analyses allowed us to confirm and further expand results from the univariate analyses, highlighting the automaticity of processing reflectional patterns in extrastriate visual areas, the MVPA analysis technique also provides the advantage of investigating whether we can differentiate between regularity types. Indeed, the univariate analysis reported in the previous section did not show significant differences in the mean response to reflection and rotation configurations. Here we therefore ask the following questions: do different regular patterns (i.e., reflection and rotation) result in different neural representations in extrastriate cortex, and how does attention modulate these representations? We trained an SVM classifier to best discriminate between the two classes of regular configurations, for each attentional task, respectively, and run a one-sample t-test for each task condition decoding accuracy against chance level, for each ROI separately. Interestingly, this analysis revealed that, when attending to regularity, our classifier was able to reliably differentiate between reflections and rotations across all ROIs, with exception of ventral areas VO1 and VO2. Attending to the luminance of the dots (a visual feature that would not require processing of the configuration of the dot patterns), resulted in similar outcomes: all ROIs but V1 and VO2 showed significantly above chance classification between reflection and rotation (Tale 2 - columns 5 and 6; Figure 4C).

Finally, given that one of the manipulations of our stimuli consisted in change of luminance, we implemented SVMs to test whether luminance category (i.e., white, light grey, and dark grey) could be decoded across the visual hierarchy, and whether attention had an effect on these responses. We therefore ran a multi-category decoding classification (i.e., white vs light grey vs dark grey) on the response patterns while attending to luminance and while attending to regularity and run one-sample t-tests, with chance level this time being 33.33%. In Table 3 we show the results of the one-sample t-tests in each ROI. Interestingly, attending to luminance resulted in above chance decoding accuracy across the visual cortex, excluding areas VO2 and LO2 (Figure 5A). When participants were engaged in the regularity task, the classifier was not able to distinguish between representations of any luminance condition in any of the ROIs investigated (all *p*>0.082).

**Figure 5.**
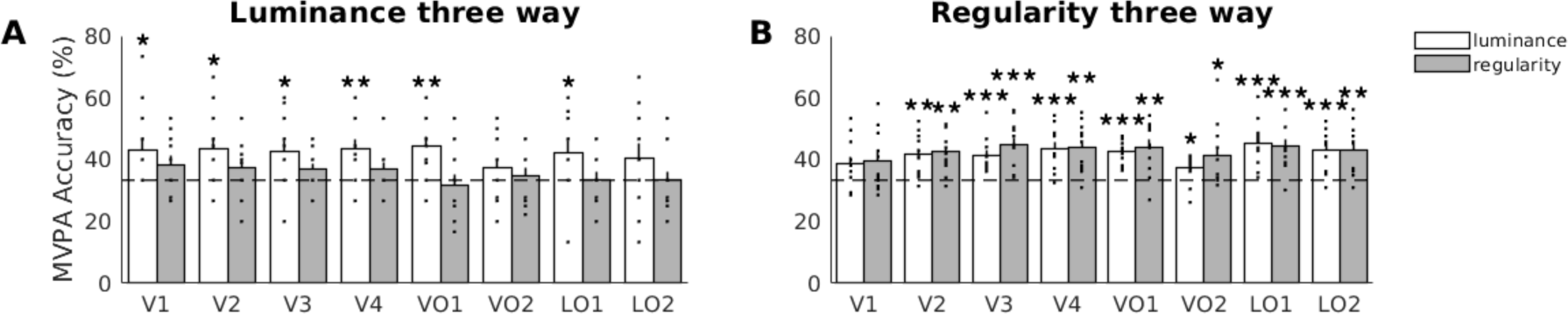
MVPA results for 3-way classification on luminance (A) and stimulus configuration (B) conditions across visual ROIs. (A) All visual areas, except VO2 and LO2, show above chance classification accuracy during the luminance (white) task, whereas attending to regularity (grey) impacts classification in all regions. (B) All visual areas, except for V1, show above chance classification accuracy during both luminance (white) and regularity (grey) tasks. Individual dot markers indicate single subject data. Dashed lines indicate MVPA accuracy at 33.33% chance. Error bars indicate +SEM. Asterisks indicate significance as follows: *<0.05, **<0.01, ***<0.001 consistent with statistics reported in Table 3 .

**Table 3.** Results of one-sample t-tests assessing classification accuracy against 33.33% chance level for multi-category (three luminance levels; three stimulus configuration levels) and task, across regions of interest. ROIs are given in each row and columns show statistical t, p-value, and Hedge’s g as a measure of effect size accounting for small sample size, for classification test (white vs light grey vs dark grey; random vs reflection vs rotation) and each task (luminance, regularity).

| ROI | Stimulus Luminance<br>(t, p, Hedge's g) |  | Stimulus Configuration<br>(t, p, Hedge's g) |  |
| --- | --- | --- | --- | --- |
|  | *luminance* | *regularity* | *luminance* | *regularity* |
| V1 | **2.335, .042, .650** | 1.609, .139, .448 | 2.155, .057, .599 | 2.140, .058, .596 |
| V2 | **2.705, .022, .753** | 1.523, .159, .424 | **3.996, .003, 1.112** | **4.465, .001, 1.242** |
| V3 | **2.631, .025, .732** | 1.929, .083, .537 | **4.871, .001, 1.355** | **4.931, .001, 1.372** |
| V4 | **3.271, .008, .910** | 1.605, .140, .447 | **4.826, .001, 1.343** | **4.415, .001, 1.228** |
| VO1 | **3.910, .003, 1.088** | -.462, .654, -.128 | **7.595, <.001, 2.113** | **4.146, .001, 1.153** |
| VO2 | 1.250, .240, .348 | .499, .629, .139 | **2.759, .020, .768** | **2.635, .025, .733** |
| LO1 | **2.235, .049, .622** | .041, .968, .012 | **5.250, <.001, 1.461** | **4.943, .001, 1.375** |
| LO2 | 1.482, .169, .412 | -.016, .987, -.005 | **4.752, .001, 1.322** | **3.849, .003, 1.071** |

The same multi-category decoding classification applied to the stimulus configuration (i.e., random vs reflection vs rotation) showed a very different pattern of results. Here, one-sample t-tests against chance level (33.33%) revealed above chance decoding accuracy beyond area V1 (all *p*<0.025) both when participants were performing the luminance and the regularity task (Figure 5B). This result aligns with the findings that information of spatial configurations is retained during sensory processing in a more automatic/task-independent fashion.

## Discussion

We presented participants with two categories of regular dot patterns, as well as random configurations, while they engaged in either a regularity-relevant (1-back task on the category of the patterns) or regularity-irrelevant (1-back task on the luminance of the dots in the patterns) task. This design allows us to tap into the mechanisms underlying symmetry processing as a proxy to perceptual organisation, the ability to organise sensory inputs into meaningful representations, and ask whether these mechanisms are modulated by attention. In line with previous fMRI studies (Keefe et al., 2018; Kohler et al., 2016; Sasaki et al., 2005; Tyler et al., 2005; Van Meel et al., 2019), we observe symmetry-selective responses to emerge in V3 and continue throughout the extrastriate visual cortex. This reliable evidence supports the knowledge of an extended network of visual areas contributing to processing regularity information, while the early visual cortex (V1 and V2) does not show a symmetry-specific change in haemodynamic response.

Notably, analysis of our task manipulation suggests an automaticity to the processing of visual symmetry: attending to the luminance of the patterns, a feature that does not require processing of the spatial organisation of the dots, still resulted in reliable symmetry-specific BOLD responses in extrastriate areas. This aligns well with evidence from (Sasaki et al., 2005), and (Keefe et al., 2018; Sasaki et al., 2005), both showing symmetry-selective responses even while participants were passively viewing dot patterns, without actively engaging in a regularity-relevant task. Furthermore, (Makin, Rampone, & Bertamini, 2020; Makin, Rampone, Morris, et al., 2020) also show strong evidence for the automatic processing of visual symmetry: the SPN is observed across a range of tasks, including regularity discrimination, colour judgements, oddball trials, and sound/colour congruency judgements. The brain, therefore, responds to retinal symmetry whether the task at hand actively requires perceptual organisation processing or not. It is important to notice that extraretinal symmetry, e.g., symmetry disrupted by perspective distortion or occlusion, is more fragile (Keefe et al., 2018; Makin et al., 2015; Rampone et al., 2019): symmetry-selective responses are measured only when regularity, and thus spatial organisation of the visual elements, is task relevant.

Our results here show an enhanced response when attending to regularity in ventral regions VO1 and VO2, while no such effect is shown by other extrastriate / object-selective areas. This boost is consistent with findings from (Keefe et al., 2018) showing strongest symmetry response in area VO1 (and LOB) that is further enhanced during symmetry detection task (compared to passive viewing). The difference in the extent of this attentional modulation in symmetry response, not observed across extrastriate areas in our data, could be linked to the difference in configuration of the stimuli: while both studies used salient, fourfold reflectional symmetry as one condition of interest, the size of the patterns differed considerably. Indeed, our stimuli were designed to optimise our multivariate analysis. Specifically, we chose a size that would allow processing of (global) spatial information while preventing eye movements. Furthermore, the location of individual dots were randomly selected within the aperture to control for potential confounds arising from repeated presentations of specific retinal information. Thus, this design strategy ensures that each pattern is specific to the regularity, rather than being a pattern (e.g., template / shared part as used in (Keefe et al., 2018; Van Meel et al., 2019) that would be modulated by the regularity. This clearly has methodological advantages, however it also sets a harder challenge to the classification routine, given that each pattern has its own, non-shared, configuration.

The results from the MVPA analysis largely agree with the univariate results: our classifier distinguished between neuronal representations elicited by reflectional patterns compared to those from random patterns. This was the case across extrastriate visual areas, including area V3 and V2, and classification performance was not affected by the specific stimulus property (luminance or regularity) that was task relevant at the time of encoding. This speaks in favour of the saliency and automaticity of symmetry processing across areas of the visual cortex.

Representations of rotational patterns could be distinguished from those of random ones across extrastriate areas when volunteers were engaged in the regularity task. During the luminance task, only object-selective regions, and area V2, showed above chance classification accuracy, suggesting that rotation is a type of regularity benefitting from attentional boost when consolidating its neuronal representation in visual cortex. These results extend our univariate findings by showing a potential role of area V2 in regularity processing. Interestingly, this area showed enhanced connectivity with ventral region V4 when processing symmetrical, versus asymmetrical, stimuli (Van Meel et al., 2019), suggesting a feedforward process where local, low-level features accessed in V2 are further processed into more complex properties along the visual hierarchy. Our observation that there is at least some involvement of V2 in representation of regularity is not strongly supported by other neuroimaging studies including our own that have largely been concerned with assessing univariate responses, which routinely emerge in V3 and beyond. However, studies of the macaque visual system have shown regularity-specific activation in area V2 for visual stimuli capturing structural features of naturalistic textures. (Freeman et al., 2013) compared single cell recordings from macaques V1 and V2 while presented with regular synthesised texture stimuli and matched noise patches and showed a vigorous response to regular, naturalistic textures for neurones in V2, compared to V1. The authors also measured modulation of BOLD responses in humans to the same stimuli, and reported regularity-specific responses in area V2, suggesting a sensitivity to higher-order features in this region that contribute to processing spatial properties of incoming sensory information. More recently, (Audurier et al., 2022) show symmetry-selective responses to wallpaper stimuli emerging from area V2, and extending to V3, V4, and V3A, in macaques. This study, replicating the experimental protocol by (Kohler et al., 2016) in humans and adopting it for macaques, thus suggests that macaque area V2 is sensitive to spatial structures within visual stimuli that are fed forward to more ventral areas (direct anatomical connections with V4; (Felleman et al., 1997; Gegenfurtner et al., 1997), thus contributing to perceptual organisation of higher order spatial properties. Our findings suggest that the increased sensitivity of multivariate approaches may have unmasked a role for V2 in regularity processing in humans, too.

As highlighted above, MVPA analysis provides the power to disentangle neuronal representations of different regular patterns within regions of interest. Our classifier shows that voxel patterns related to reflection can be distinguished from those related to rotational configurations in extrastriate, object-selective areas. Importantly, the decoding of spatial information here is also irrespective of the task performed: attending to a first order property of the stimulus such as luminance does not affect classification. Again, these results not only speak to the automaticity of regularity processing, but extend the univariate findings: by exploiting fine-grained spatial arrangements of multi-voxel response patterns, it is possible to gain processing information that would otherwise be averaged out in univariate analyses.

This suggests reflection and rotation do not activate the same neural populations in the extrastriate cortex - however SPN priming studies show that reflection and rotation representations are not completely independent. Prior presentation of rotation increases the SPN response to reflection and vice versa (Experiment 5 in (Makin et al., 2021).

Unexpectedly, decoding here shows above chance accuracy also for area V1 when participants are engaged in the regularity task. Early visual cortex has consistently been reported to not show sensitivity to regular structures, however, first-order information such as local orientation and spatial scale are represented in V1, and these properties could be then input to further processing in extrastriate cortex. (van der Zwan et al., 1998) induced tilt aftereffects using symmetrical dot patterns, thus treating orientation of axes-of-symmetry like orientation defined by luminance. When testing monocularly, the tilt aftereffect elicited by adapting to the orientation of the axes-of-symmetry in the pattern, did not transfer to the unstimulated eye. This suggests that orientation information of regular patterns is available in early visual cortex (V1) and is then further processed downstream.

Finally, we were able to ask our classifier to identify voxel response patterns relating to different luminance stimulus conditions. While luminance is a global property of the dot patterns, this can be resolved at the local level, i.e. attending to individual elements of the patterns will suffice to perform the luminance task here, and equally this property could be encoded when attending to the global structure of the patterns in the regularity task. Thus, one would hypothesise that first order visual properties such as luminance might be automatically accessible under different task and attentional conditions. However, our MVPA results show that this is not the case: decoding luminance information is task-selective, and this feature is available across regions of the visual cortex only when overtly attending to it. While at first counterintuitive, this finding speaks in favour of perceptual organisation; our visual system engages in perceiving and representing spatial properties in the effort of generating meaningful representations despite a large variability of first order property conditions such as luminance. The visual system, and extrastriate cortex more specifically, can therefore discount and remove representation of sensory information that are not relevant for the task at hand and do not determine an invariant representation of the emergent spatial structure. This is a feature known in object-selective areas such as the LOC (Kourtzi & Kanwisher, 2001), with LO1 and LO2 regions shown to independently process spatial aspects of complex objects - orientation and shape, respectively (Kourtzi & Kanwisher, 2001; Silson et al., 2013).

## Conclusion

In summary, we show symmetry-specific responses to emerge from area V3 and extend throughout extrastriate/ventral and object-selective visual areas. Multi-voxel pattern analysis supports this finding and suggests a role in processing of spatial information already in area V2. Furthermore, the contrast in the task effect of the classification performance for distinguishing regularity and luminance is intriguing. The fact that the dot luminance could not be classified when participants attend to regularity indicates that this first order visual property is likely discounted from the neural representation in a task specific way. In contrast, the spatial properties of the stimuli can be decoded even when participants did not attend to those features. We propose therefore that the visual system, most particularly the extrastriate visual areas, computes and represents these global properties largely automatically and underpins perceptual organisation.

## Acknowledgements & Conflicts of Interest

We thank the staff at the York Neuroimaging Centre for support with MRI scanning. We have no conflicts to declare.

## Open Research - Data availability statement

The summary data and code that support the findings of this study are openly available via Open Science Framework at https://osf.io/t7esz/?view_only=132bc7e6178741f4b2b10c045c355fcb. Raw and processed fMRI data will be openly available via OpenNeuro soon (DOI: 10.18112/openneuro.ds004715.v.1.0.2).

